# Identification of herboxidiene features that mediate conformation-dependent SF3B1 interactions to inhibit splicing

**DOI:** 10.1101/2020.11.18.387712

**Authors:** Adriana Gamboa Lopez, Srinivasa Rao Allu, Patricia Mendez, Guddeti Chandrashekar Reddy, Hannah M. Maul-Newby, Arun K. Ghosh, Melissa S. Jurica

**Affiliations:** Department of Molecular Cell and Developmental Biology, University of California, Santa Cruz, CA, USA; Center for Molecular Biology of RNA, University of California, Santa Cruz, CA, USA; Department of Chemistry and Department of Medicinal Chemistry, Purdue University, West Lafayette, IN, USA

**Keywords:** SF3B, pre-mRNA splicing, SF3B inhibitors, herboxidiene analogs

## Abstract

Small molecules that target the spliceosome SF3B complex are potent inhibitors of cancer cell growth. The compounds affect an early stage of spliceosome assembly when U2 snRNP first engages the branch point sequence of an intron. Recent cryo-EM models of U2 snRNP before and after intron recognition suggest several large-scale rearrangements of RNA and protein interactions involving SF3B. Employing an inactive herboxidiene analog as a competitor with SF3B inhibitors, we present evidence for multiple conformations of SF3B in the U2 snRNP, only some of which are available for productive inhibitor interactions. We propose that both thermodynamics and an ATP-binding event promote the conformation conducive to SF3B inhibitor interactions. However, SF3B inhibitors do not impact an ATP-dependent rearrangement in U2 snRNP that exposes the branch binding sequence for base pairing. We also report extended structure activity relationship analysis of herboxidiene, which identified features of the tetrahydropyran ring that mediate its interactions with SF3B and its ability to interfere with splicing. In combination with structural models of SF3B interactions with inhibitors, our data leads us to extend the model for early spliceosome assembly and inhibitor mechanism. We postulate that interactions between a carboxylic acid substituent of herboxidiene and positively charged SF3B1 sidechains in the inhibitor binding channel are required to maintain inhibitor occupancy and counteract the SF3B transition to a closed state that is promoted by stable U2 snRNA interactions with the intron.

## INTRODUCTION

The spliceosome is responsible for removing introns from gene transcripts in eukaryotes to generate messenger RNAs (mRNA). It assembles on each intron to be removed from a transcript from five U-rich small nuclear RNAs complexed in ribonucleoproteins (snRNPs), which join with dozens of additional proteins through a complicated series of interactions and structural rearrangements. In early spliceosome assembly, U2 snRNP, which consists of U2 snRNA, core proteins and the SF3A and SF3B multi-protein complexes, joins the intron to form the A-complex spliceosome. U2 snRNA base pairs with the neighboring nucleotides of the branch point sequence (BPS) to specify an adenosine residue near the end of intron as the branch point ^1,2^. The adenosine is eventually positioned by U2 snRNA in the spliceosome’s catalytic center to participate in the first chemical step of splicing ^3–5^. Selection of the BPS also designates the 3’ splice site at the end of intron as the first downstream AG dinucleotide. Additional proteins contribute to BPS selection, such as the U2 auxiliary factor (U2AF) which recognize the downstream polypyrimidine tract ^6^. Notably, several proteins involved in BPS selection, including SF3B1 and U2AF1, are frequently mutated in some cancers and dysplasia, which suggests a role for intron definition in carcinogenesis ^7–9^. Specific SF3B1 cancer mutations are linked to use of an aberrant branch point and 3’ splice site in some transcripts, indicating that SF3B1 normally helps ensure fidelity of BPS selection ^10–12^.

Recent cryo-EM structural models of the spliceosome captured in the early stages of assembly provide some clues for how SF3B1 participates in BPS recognition ^1,13–16^. The N- and C-terminal regions of the central C-shaped HEAT-repeat domain of SF3B1 come together and appear to clamp onto the branch helix formed by U2 snRNA and the intron. This conformation sequesters the bulged branch point adenosine into a pocket formed between PH5FA and SF3B1 HEAT repeats 15 and 16. Interestingly, in structures of U2 snRNP and isolated SF3B protein complex, the same domain exhibits a more open conformation that hinges at the same two HEAT repeats ^17,18^. Together, these structures imply that closing the SF3B clamp is a critical step in BPS selection.

Further support for this model originates with the small molecule splicing inhibitors pladienolide B (PB), spliceostatin A (SSA) and herboxidiene. These natural products derived compounds were originally identified as potent inhibitors of tumor cell growth, and later shown to target SF3B and interfere with spliceosome assembly when U2 snRNP joins the intron to form the spliceosome A-complex ^19–24^. Structures of SF3B bound to PB revealed the inhibitor cradled in a channel between PHF5A and SF3B1 HEAT repeats 15 and 16 in the open conformation ^25,26^. Point mutations in either SFB1 or PHF5A residues in the channel render cells resistant to SF3B inhibitors, which confirms the channel as the inhibitor target ^27,28^. SF3B inhibitors are proposed to hold SF3B1 in an open conformation. In structures where SF3B1 is closed over the branch helix, many residues contacting the inhibitor in the channel are moved to form the pocket for the branch point adenosine. Therefore, even though the residues are situated in a different position when SF3B1 is closed, SF3B inhibitors effectively compete with the adenosine for the same interactions.

We previously reported that splicing inhibition by PB, SSA and herboxidiene can be masked by addition of an excess of inactive analogs of each compound ^29^. Furthermore, inactive analogs of one structural family rescued splicing from inhibitors of the other two families. We hypothesized that the inactive analogs compete with the SF3B inhibitors for the same binding site, but lack a chemical feature that is required either for more stable binding or for interfering with SF3B function. In this study, we test this hypothesis and investigate factors that influence splicing inhibition. Our data show that relative to an inactive herboxidiene analog, inhibitors exchange from SF3B more slowly. We present evidence that SF3B conformation controls access to the inhibitor binding channel, and is regulated by an ATP-binding event. Testing a series of herboxidiene analogs for their ability to inhibit splicing or to compete with active inhibitors, we conclude that SF3B1 interactions with the C1 carboxylic acid of herboxidiene are essential for the inhibitor to maintain binding in the context of SF3B closure over the branch helix.

By interrogating chemical features of herboxidiene, our results are important for its potential development as a chemotherapeutic. Some SF3B inhibitor derivatives have already shown promise in this regard. For example, H3B-8800 decreases the volume of human tumors implanted in mice regardless of SF3B1 mutation status ^30^. However, another PB derivative failed to pass Phase I clinical trials ^31^. SSA derivatives are also being pursued as cancer treatments. Recently, Yoshikawa et. al. 2020 reported a new SSA derivative that represses expression of a chemotherapeutic-resistant splicing variant and exhibits low toxicity mice ^32^. Clearly further investigation into the mechanism of SF3B inhibitors is needed to realize their promise for human health. Herboxidiene represents a promising platform for creating new drug candidates, because it has a simpler structure that allows us to meet the synthetic challenges required for detailed structure activity relationship (SAR) studies that are crucial for tuning potency and specificity.

## RESULTS

### Order of addition affects competition between SF3B inhibitors and inactive analogs

We previously showed that when 1 μM herboxidiene, PB or SSA alone is included in an *in vitro* splicing reaction we do not detect splicing products. However, when 100 μM of an inactive analog of herboxidiene (iHB) is added simultaneously, splicing products are produced at levels nearly equivalent to control reactions with no added inhibitor. We tested whether the order of addition affects competition between SF3B inhibitors and iHB competitor. We reasoned that if inhibitor interactions exchange rapidly with SF3B, then incubating the inhibitor in nuclear extract prior to adding excess inactive competitor would have no effect on the competitor’s ability to rescue *in vitro* splicing from inhibition. Alternatively, if the inhibitor exchanges slowly or irreversibly interferes with SF3B function, then excess inactive competitor added after the inhibitor would not be able to replace it and result in low splicing efficiency. To distinguish between these two possibilities, we incubated 1 μM PB in nuclear extract for 10 minutes, and then added 100 μM iHB for an additional 10 minutes. As controls, we replaced both compounds with DMSO, and reversed the order of addition. We then tested the nuclear extracts for *in vitro* splicing efficiency. As expected, nuclear extract treated with PB or SSA followed by DMSO yielded no splicing products (Figure1a lanes 3 and 6, and 1B). An excess of inactive competitor iHB added before inhibitor PB or SSA rescues splicing efficiency to the level of DMSO controls, consistent with previous results with simultaneous addition (Figure 1a lanes 5 and 8, and 1B) ^29^. In contrast, when PB or SSA is incubated with nuclear extracts before addition of excess iHB competitor, splicing efficiency is greatly reduced, although not completely lost relative to the inhibitor alone (Figure 1a, lanes 3-5 and 6-8, and 1B). We also tested herboxidiene and less potent inactive competitor analogs of PB and SSA in the same manner with similar outcome (Supplemental Figure 1). Although our assay does not directly measure binding, these results are consistent with a very slow off-rate for the inhibitors. Alternatively, they may reflect extended inactivation of SF3B by an unknown mechanism. In either case, the result opens the door to using the competition assay to assess factors that affect the interaction between SF3B and its inhibitors.

**Figure 1.**
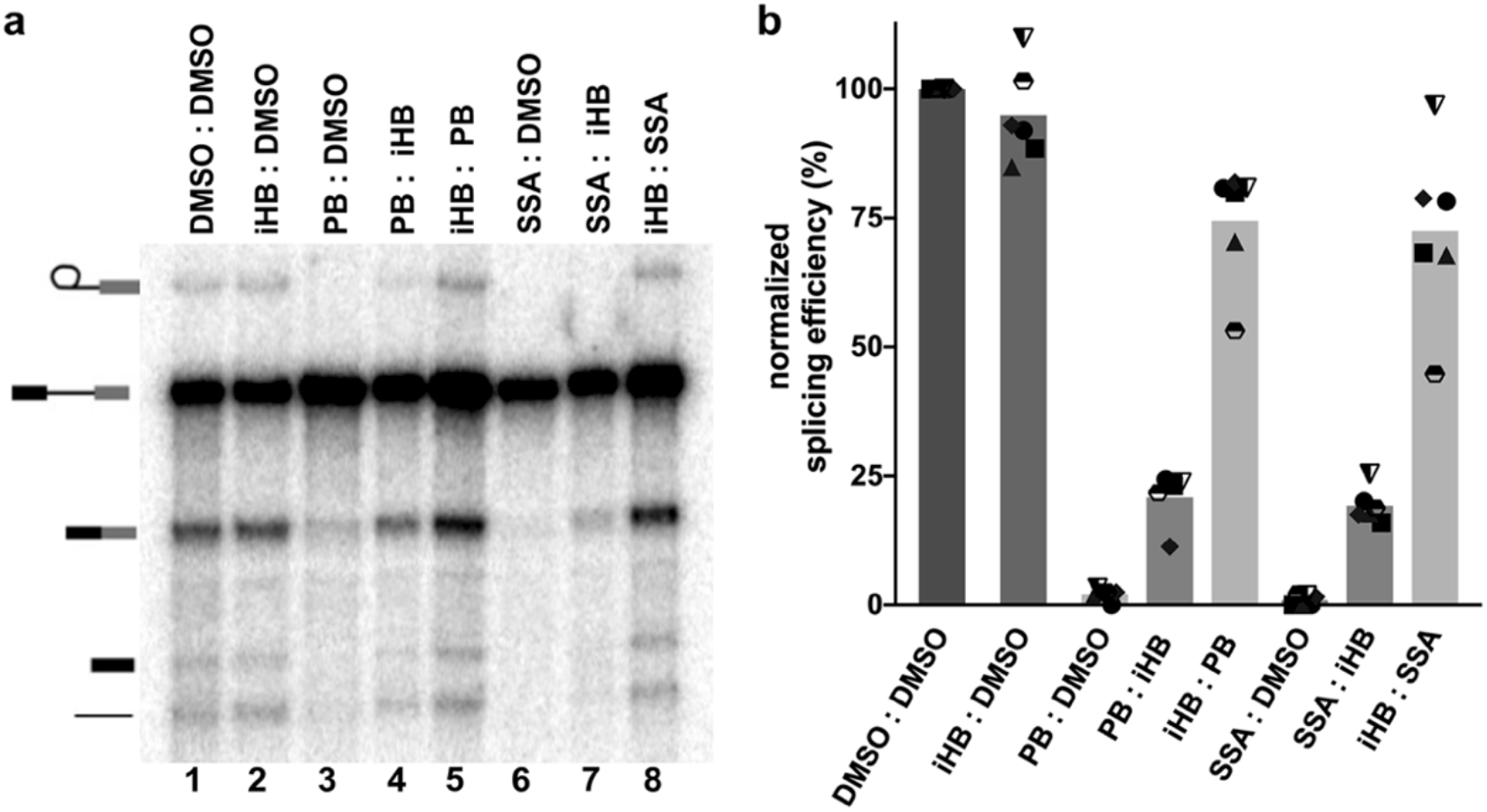
Order of addition affects competition between SF3B inhibitors and an inactive herboxidiene analog. (a) Representative denaturing gel analysis of *in vitro* splicing. Prior to addition of splicing substrate, nuclear extracts were first incubated at 4°C for 10 minutes with one of the following compounds: DMSO, iHB (100 μM), PB (1 μM) or SSA (1 μM) followed by DMSO, SSA (1μM), PB (1μM) or iHB (100μM) for another 10 minutes, as indicated. Band identities are illustrated on the left as (top to bottom): lariat intron product, pre-mRNA substrate, mRNA product, 5’ exon intermediate and linear intron product. (b) Splicing efficiency quantified from denaturing gels plotted relative to DMSO control. Order of addition is denoted as **1st compound: 2nd compound**. Mean normalized splicing efficiency is displayed as bars, with experimental replicates values represented by different shape.

### Inhibitor interactions with SF3B are modulated by temperature

Because structural models imply that an open SF3B1 conformation is required for inhibitor binding^1,26^, we reasoned that temperature may modulate SF3B conformational dynamics and affect accessibility of the inhibitor channel. We repeated the same order of addition assay and compared the effect of incubating inhibitors at 4°C vs. 30°C (Figure 2a). In the absence of competition, incubation temperature has no significant effect on splicing or splicing inhibition (Figure 2b, c and Supplemental Figure 2a, b). The result is different in the context of competition. When SSA, PB or herboxidiene is first incubated in nuclear extract at 30°C before addition of excess iHB competitor, the extract’s splicing efficiency is lower compared to when they are first incubated at 4°C (Figure 2b lanes 6 vs 7, 8 vs 9 and 10 vs 11). The effect of a higher temperature incubation providing inhibitors a stronger competitive advantage relative to a lower temperature incubation also holds for less potent inactive competitor analogs of PB and SSA (Supplemental Figure 2b). The difference is consistent with differential accessibility of the inhibitor binding channel in SF3B depending on its conformation. We speculate that higher temperature favors the SF3B1 open conformation to give the inhibitor more opportunity to interact with SF3B before the competitor is added, which enables the inhibitor to prevent splicing when substrate is added.

**Figure 2.**
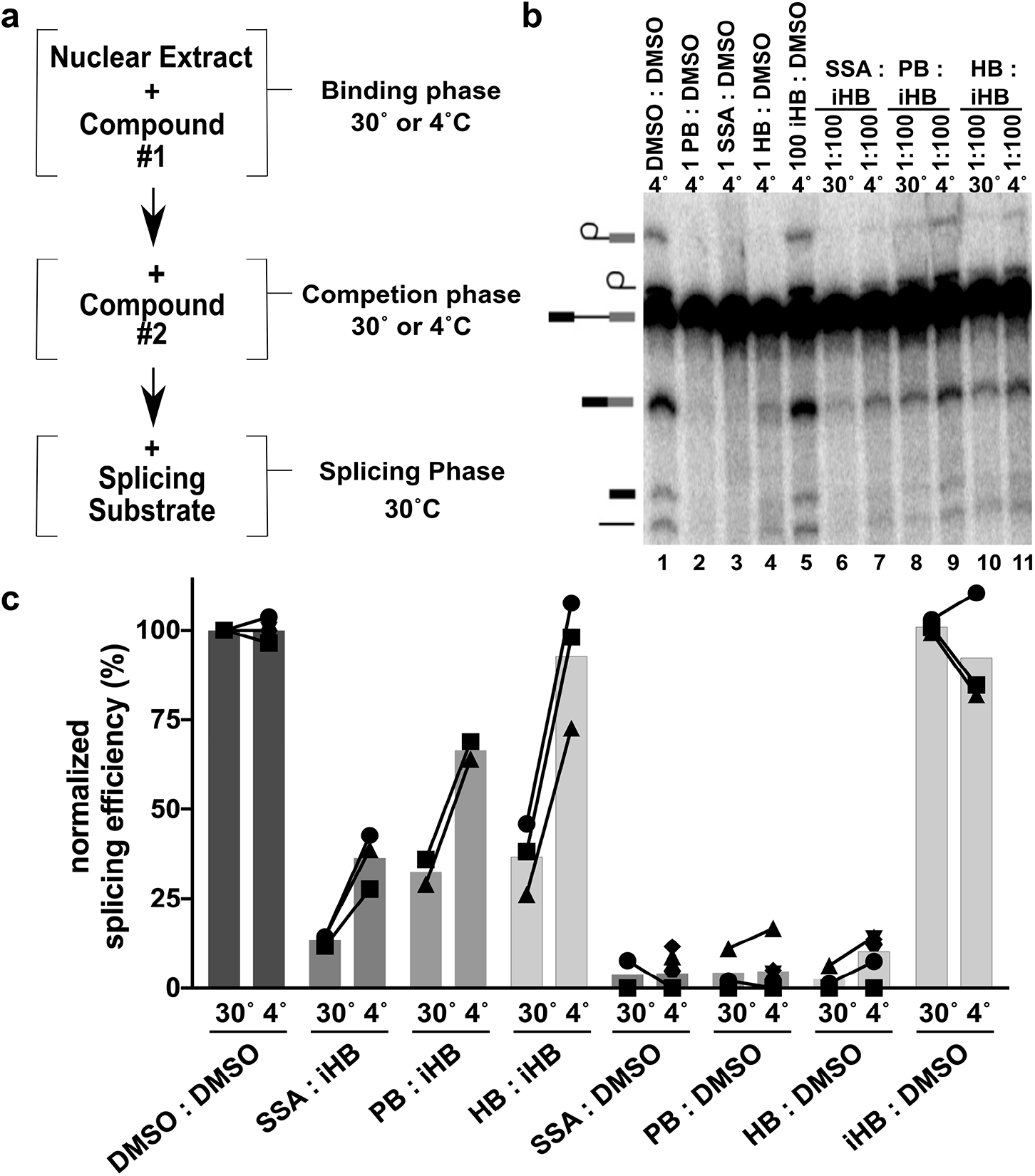
Temperature affects competition between SF3B inhibitors and inactive analogs. (a) Schematic of order of addition assay. (b) Representative denaturing gel analysis of *in vitro* splicing. Prior to addition of splicing substrate, nuclear extracts were first incubated 10 minutes at 4°c or 30°c with one of the following compounds: DMSO, SSA (1 μM), PB (1μM), HB (1μM) or iHB (100 μM), followed by DMSO or iHB (100μM) for another 10 minutes. Band identities are illustrated on the left as (from top to bottom) lariat intermediate, lariat intron product, pre-mRNA substrate, mRNA product, 5’ exon intermediate and linear intron product. (c) Splicing efficiency quantified from denaturing gels plotted relative to DMSO control. Order of addition is denoted as **1st compound: 2nd compound**. Mean normalized splicing efficiency is displayed as bars, with experimental replicates values represented by different shapes and linked to highlight differences in splicing rescue. Representative denaturing gel for control reactions with SSA, PB and HB (1 μM) or iSSA, iPB and iHB (100 μM) followed by DMSO is shown in Supplementary Figure 2a.

### Temperature-dependent interactions with SF3B are not modulated by ATP hydrolysis

The mechanisms controlling SF3B conformation are not fully understood. A recent cryo-EM structure of the U2 snRNP revealed SF3B in the open conformation with the inhibitor binding channel accessible ^18^. Notably, the RNA-dependent ATPase DDX46, the human ortholog of yeast Prp5, is positioned close to SF3B1 with an extended N-terminal helix making direct contact with HEAT repeats that must move with closing. Additionally, the protein HTATSF1 contacts both sides of the SF3B1 clamp, suggesting that it may serve as a brace for the open conformation. In addition to the structures, genetic and biochemical evidence show that both proteins are displaced prior to completion of A-complex assembly^16,33,34^. It is possible that either the positioning or removal of DDX46 and HTATSF1, and thus SF3B1 conformation, are dependent on ATP hydrolysis. In the competition assays described above, we included ATP during inhibitor incubation in the nuclear extracts. To test the hypothesis that the temperature dependent effects on inhibitor interactions with SF3B were mediated by an ATP hydrolysis event, we repeated the splicing assay but excluded ATP during the competition phase of the experiment. Surprisingly the absence of ATP during the competition phase did not abolish the effect of temperature on splicing inhibition. ATP-depleted extracts incubated with inhibitor at 30°C followed by excess competitor still exhibited lower splicing efficiency relative to competition at 4°C (Figure 3a lanes 4 vs. 5, 6 vs. 7, and 3b). We conclude that SF3B1’s ability to transition from a closed conformation that is refractory to inhibitor interaction to an open conformation that SF3B inhibitors can access is thermodynamically influenced. Still, the presence of ATP does favor inhibitor interactions at both 4°C and 30°C (Figure 3a lanes 4 vs. 6, 5 vs. 7, and 3b), indicating that ATP promotes inhibitor access to SF3B in a temperature-independent manner. Because most ATPases are inactive at 4°C, the difference may reflect ATP-binding, rather than hydrolysis, stabilizes the open conformation of SF3B, possibly by DDX46.

**Figure 3.**
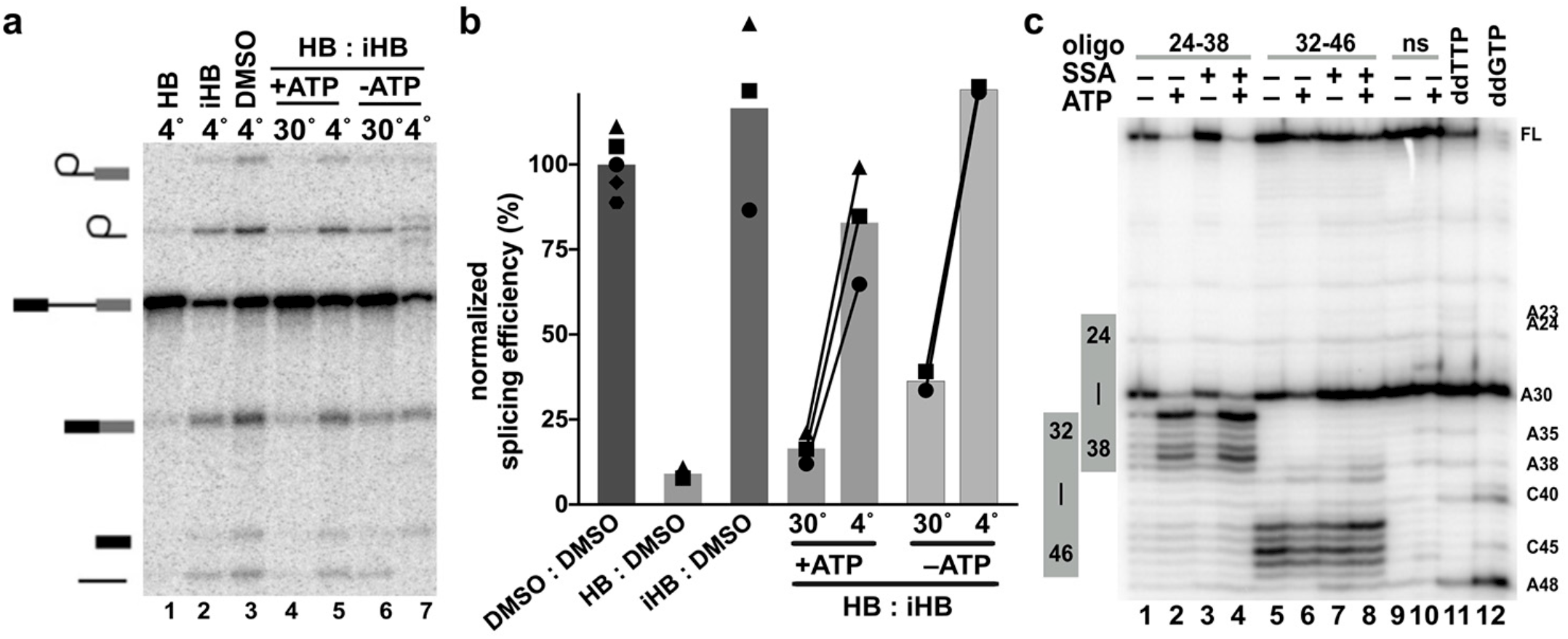
Temperature-dependent modulation of SF3B inhibition is independent of ATP. (a) Representative denaturing gel analysis of *in vitro* splicing. Prior to addition of splicing substrate, nuclear extracts were first incubated 10 minutes at 4°C or 30°C with or without ATP and one of the following compounds: DMSO, HB (1μM) or iHB (100 μM) followed by DMSO or iHB (100μM) for another 10 minutes. Band identities are illustrated on the left as (from top to bottom) lariat intermediate, lariat intron product, pre-mRNA substrate, mRNA product, 5’ exon intermediate and linear intron product. (b) Splicing efficiency quantified from denaturing gels plotted relative to DMSO control. Order of addition is denoted as **1st compound: 2nd compound**. Mean normalized splicing efficiency is displayed as bars, with experimental replicates values represented by different shapes and linked to highlight differences in splicing rescue. (c) Primer extension analysis of targeted DNA-oligo mediated RNase H digestion of U2 snRNA in nuclear extracts. Extracts were incubated with and without ATP in the presence and absence of 1 μM SSA at 30°C for 10 minutes prior to addition of oligos complementary to the indicated nucleotides. (ns = non-specific oligo, ddTTP and ddGTP = sequencing lanes)

### ATP-dependent changes in U2 snRNA BBS accessibility are not influenced by SF3B1 inhibitors

Stable incorporation of U2 snRNP in A-complex to establish the branch helix requires at least one ATP dependent step ^35–37^. ATP is also required for U2 snRNP association with pre-mRNA in the presence of both SSA and PB to form an unstable A-like complex in which the branch helix is not fully formed^21,22,38^. U2 snRNP conformation is also regulated by ATP^23,39,40^. Folco *et al* reported that the PB analog E7107 inhibits U2 snRNPs ability to engage an oligo containing the branch sequence, but that inhibition is bypassed by an ATP-dependent rearrangement of the snRNP ^23^. We hypothesized that the small improvement of inhibitor competition conferred by the presence of ATP at both 4°C and 30°C is due to the same ATP-dependent rearrangement. To test this idea, we examined U2 snRNA accessibility in the presence and absence of ATP and inhibitor.

Endogenous RNase H in nuclear extract cleaves RNA in the RNA/DNA hybrids that form upon addition of DNA oligos that are complementary to regions accessible for base pairing. We tested DNA oligos targeting U2 snRNA nucleotides between Stem I and Stem IIa, and mapped cleavage sites using primer extension. As predicted by previous studies, cleavage efficiency changes in some regions when extracts are first incubated with ATP^40^. The most predominate change is an increase in cleavage at nucleotides 33 through 38 with an oligo complementary to nucleotides 24-38 (Figure 3c lanes 1-4). Notably, these nucleotides are the GUAGUA nucleotides of the branch binding sequence that forms base pairs with an intron. They also sit in the loop region of the branchpoint-interacting stem loop (BSL) as recently visualized in the U2 snRNP structure^18^. Because ATP increases RNA cleavage in this region, an unwinding of the BSL, potentially by DDX46, is a possible explanation for the increased accessibility. Alternatively, release of DDX46 from the snRNP may allow access to the DNA oligo probe. Surprisingly, the SF3B inhibitor SSA does not interfere with the ATP-dependent changes in cleavage (Figure 3c). It is therefore possible that exposing the branch binding sequence mediates the ATP-requirement for U2 snRNP associates with an intron in the presence of inhibitor. The result also suggests that this event is independent of SF3B conformation. Furthermore, the previously described ATP-dependent conformational change in U2 snRNP inhibited by SF3B remains to be characterized.^23^

### Inactive analogs of SF3B inhibitors only lose some their competitive advantage over time

The idea that splicing inhibitors preventing SF3B closure raises an interesting question: How does iHB block SF3B inhibitor action, presumably by competing for the SF3B channel, while at the same time not affect splicing? One possible explanation is that the inactive compound exchanges rapidly from the binding site in comparison with the apparent slow off-rate for active compounds suggested by our order of addition experiments. If true, when inactive compound is incubated in extract before adding inhibitor, then competition will not show the same temperature dependence, because the inactive compound be quickly replaced by an active inhibitor with the final shift to 30°C for the splicing reaction. Furthermore, the ratio of active to inactive compound should affect both the extent and rate of competition. To test these predictions, we repeated the order of addition experiment at both 4° and 30°C, but added 100μM, 10μM or 1μM iHB first for 10 minutes, and followed with 1 μM SSA for another 10 minutes. As predicted, splicing is rescued to the same level at both temperatures when an excess of inactive competitor is added first (Figure 4a lanes 2 vs. 3,6 vs.7,9 vs. 10 and 4b), and holds for competitions with inactive analogs of PB and SSA followed by herboxidiene, PB or SSA (Supplemental Figure 3a, b, c). When at a 1:1 ratio splicing was largely inhibited, but the small amount of splicing rescue was slightly higher for competition at 4°C relative to 30°C. This result is consistent with SSA having increased access to SF3B at the higher temperature as the inactive competitor leaves.

**Figure 4.**
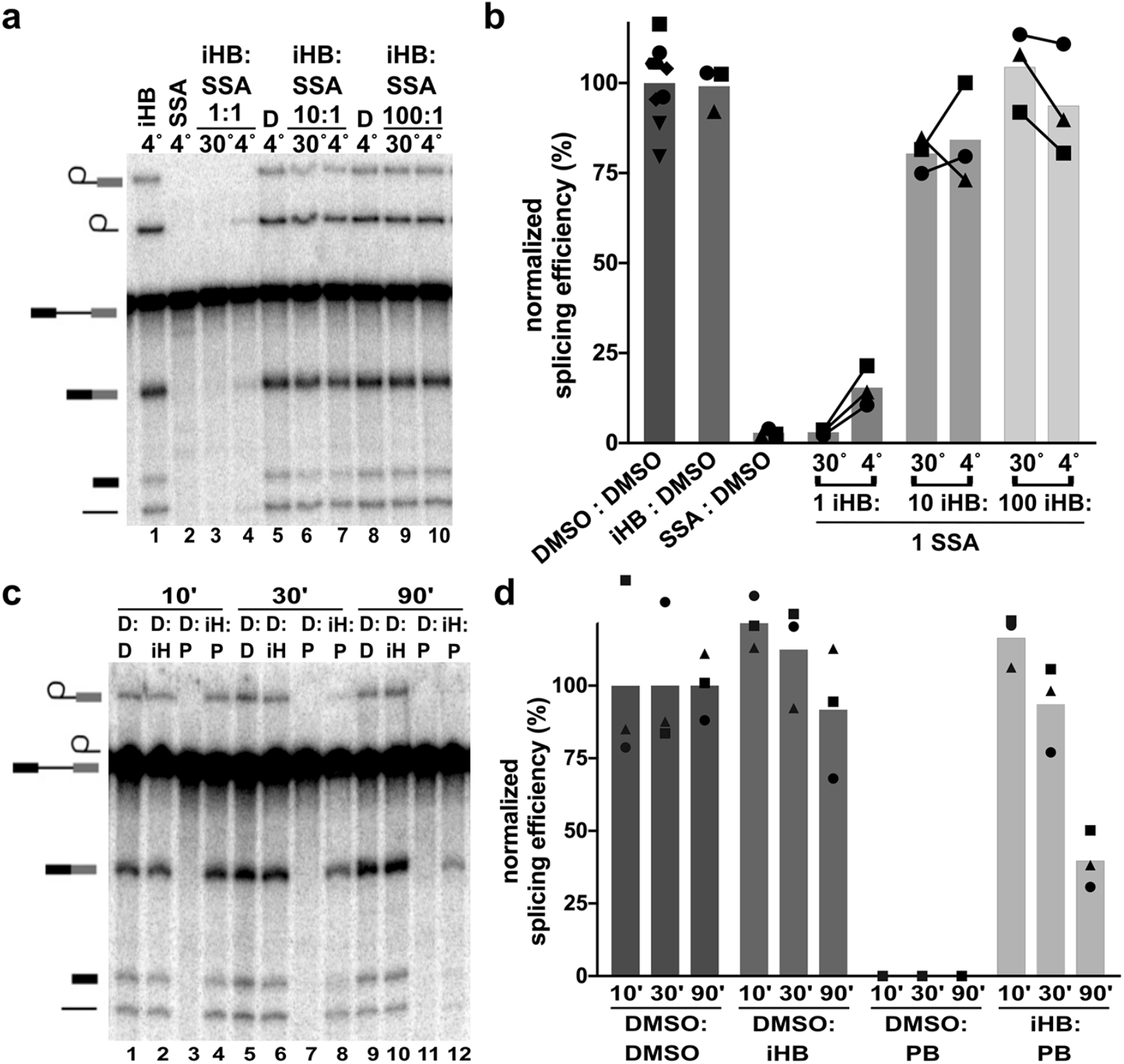
SF3B inhibitors exchange more slowly than inactive analogs. (a) Representative denaturing gel analysis of *in vitro* splicing. Prior to addition of splicing substrate, nuclear extracts were first incubated 10 minutes at 4°C or 30°C with or without ATP and one of the following compounds: DMSO control, iHB (1, 10 or 100 μM) or SSA (1 μM) followed by DMSO or SSA (1 μM) for another 10 minutes. Band identities are illustrated on the left as (from top to bottom) lariat intermediate, lariat intron product, pre-mRNA substrate, mRNA product, 5’ exon intermediate and linear intron product. (b) Splicing efficiency quantified from denaturing gels as shown in panel (a) plotted relative to DMSO control. Order of addition is denoted as **1st compound: 2nd compound**. Mean normalized splicing efficiency is displayed as bars, with experimental replicates values represented by different shapes and linked to highlight differences in splicing rescue. (C) Representative denaturing gel of *in vitro* splicing as described for panel (a), except the second incubation was carried out for 10, 30 or 90 minutes and all incubations were at 30°C. (D= DMSO, iH = iHB, P=PB) (D) Quantification of *in vitro* splicing assays similar to panel B.

Differences in exchange rates for inactive competitors vs. SF3B inhibitors also predict that extending competition for a longer period of time should result in decreased splicing rescue when inactive competitor is followed by an inhibitor that has much slower exchange rate. We tested this expectation by incubating iHB in nuclear extract for 10 minutes at 30°C and then adding PB to compete for 10, 30 or 90 minutes (Figure 4c lanes 1-4, 5-8, 9-12 respectively and 4d). To further bias competition toward the active inhibitor, we used a ratio of 10:1 iHB to PB. Although splicing efficiency decreases somewhat with longer competition, iHB’s competitive advantage still remains after 90 minutes with only a 50% reduction relative to shorter competitions of 10 or 30 minutes (Figure 4c, d). Coupled with its ability to also rescue splicing at relatively low concentrations (1:1 and 1:10), it appears that even though iHB’s exchange rate from SF3B is higher than active inhibitors, it not rapid enough to explain why iHB has no effect on splicing.

### Chemical features of herboxidiene differentially impact splicing inhibition and competition

To further explore the chemical features of herboxidiene (**1**) important for splicing inhibition and competition we tested a series of analogs in which we varied the substituents at C1 and C6 positions (see Figure 5 for compound modifications numbered (**1**)-(**20**). The inactive herboxidiene analog iHB (**14**) that we used for our competition assays differs from the active parent compound herboxidiene at two positions: an additional hydroxyl group at C5 of the tetrahydropyran ring, and conversion of the C1 carboxylic acid to its methyl ester. The C1 methyl ester is likely responsible for the lack of splicing inhibition, because the presence of the C5 hydroxyl group alone (**2**) has a minimal effect on compound activity ^41^. The parent compound, herboxidiene (**1**), has an IC_50_ value for *in vitro* splicing of 0.3 μM ^41^. Removing the methyl group at C6 (**3**) or replacement with a methylene group (**4**) had essentially no effect on compound activity, as their IC_50_ values remain sub-micromolar comparable to herboxidiene (**1**). Flipping the chirality of the methyl group (**7**), creating a di-methyl (**6**) or cyclopropane at C6 (**8**) each resulted in >4-fold decreased inhibitory activity, while addition of a methyl ether (**9**) dropped activity >10-fold. Inclusion of an epoxy group (**17**and **18**) or a dichlorocyclopropane functionality (**15**) resulted over 500-fold decreased activity with an IC_50_ value greater than 100 μM, which we classify as inactive. From these results, we conclude that the presence and orientation of the methyl group at C6 in herboxidiene is not crucial for the inhibitor’s activity. However, increasing the size of the group at that position does have a deleterious effect. We also tested some of alterations at C6 (dimethyl, cyclopropane, methylether, dichlorocyclopropane) in combination with a methyl ester at C1. In this context, all compounds showed no activity at 100 μM, further supporting a critical role for the carboxylic acid functionality at C1. Notably, substitution of the carboxylic acid at C1 with carboxamide (**10**) along with an additional hydroxyl at C5 (**12**) strongly reduced splicing inhibition (>10-100-fold), but did not completely inactivate the compound in our assay system. Converting the C1 carboxylic acid to its hydroxymethyl derivative (**11**), in the context of the normally benign C5 hydroxyl addition, also strongly affected activity.

**Figure 5.**
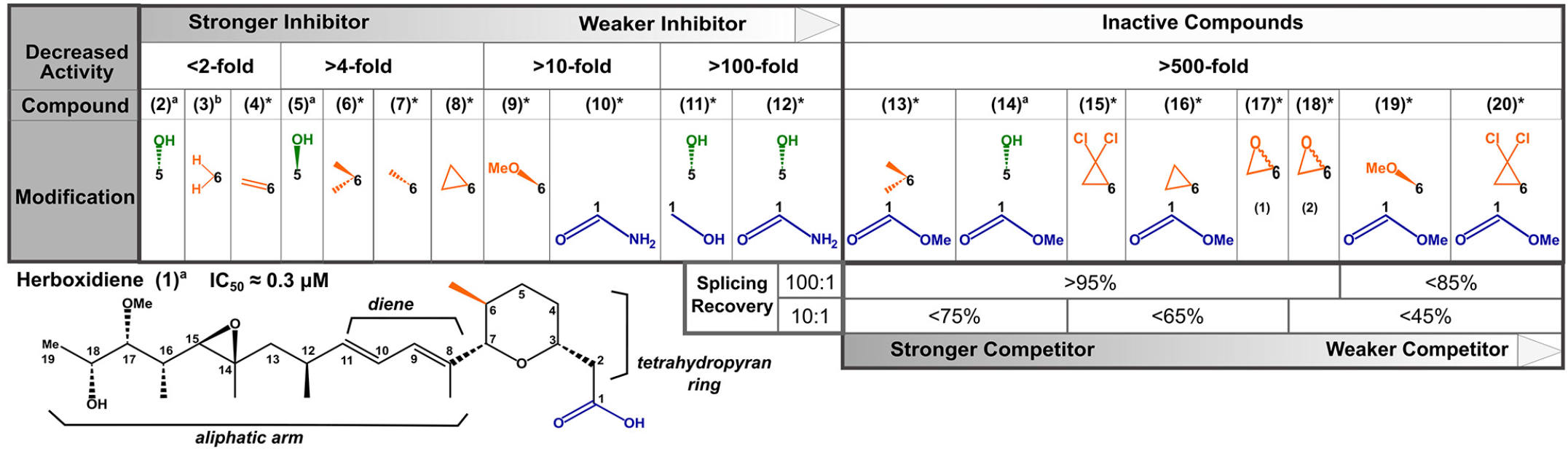
Structure activity relationships for herboxidiene activity and SF3B interaction. Summary of SAR results relative to the chemical structure of herboxidiene with modifications colored by their location and grouped by the magnitude of effect on inhibitor activity relative to the parent compound. Compounds with >500-fold decrease in inhibitory activity were tested for their ability to rescue splicing activity with 1 μM active splicing inhibitor at 100-fold and 10-fold excess. * indicates compounds new to this study, a^41^, b^46^.

We next asked at what level inactive herboxidiene analogs are able to compete with an inhibitor. In all cases, addition of most inactive analogs at 100 μM rescued >95% of splicing activity normally completely inhibited by 1 μM PB, although two compounds with both a methyl ester group at C1 and a bulky adducts at C6, rescue was slightly reduced. At a 10:1 ratio, compounds with larger adducts at C6 (compounds **15-20**) again yielded lower splicing rescue. We conclude that both these positions contribute to herboxidiene’s ability to inhibit splicing, although the functionality at C1 affects the ability to interfere with SF3B function more that it affects its ability to initially occupy the inhibitor binding channel.

## DISCUSSION

Herboxidiene, like PB and SSA, is a potent inhibitor of splicing. We identified modifications to herboxidiene that render it inactive as an inhibitor, but is nevertheless able to compete quite well with active inhibitors. Further structure-activity studies indicate that positions C1 and C6 in the tetrahydropyran moiety contribute to both herboxidiene’s interaction with SF3B. When modeled into the PB binding site of the SF3B crystal structure, the groups at these two positions are predicted to interact with residues from the U2 snRNP proteins PHF5A and SF3B1 (Figure 6a). The methyl group at C6 is close to a surface of PHF5A that remains relatively statice in the context of SF3B1 conformational changes, and larger adducts could sterically clash, which may explain why compounds with these adducts (for example, **15**) exhibit both decreased potency as splicing inhibitors and decreased ability to compete with active inhibitors. On the other hand, the carboxyl at group at C1 in interacts with residues in HEAT repeat 15 of SF3B1, which undergo a large structural shift when SF3B1 closes to engage the branchpoint adenosine of an intron. Modifications at C1 decrease herboxidiene’s activity, showing that the position is important for inhibition, but have less impact on competition. Extended competitions in which inactive competitor iHB is added first are consistent with iHB (**14)**having a higher exchange rate from the inhibitor binding channel relative to strong inhibitors. Still, why the inactive compounds are able to block SF3B inhibitors for extended periods of time, but do not block engagement of the intron at the same level is puzzling. It may be that our changes at C1, which removes a potential negative charge, do not affect affinity for the inhibitor binding channel present in an open SF3B1. However, those changes may no long interact with and potentially stabilize the nearby positively charged side chains that rearrange to contact the intron during SF3B1 closing. Consistent with this idea, SF3B1 mutants have been identified as resistant to PB and herboxidiene ^27,42^, and these mutations map to the positively charged amino acids (K1070, K1071, R1074, R1075) that are situated adjacent to C1 in our model (Figure 6b). We speculate that inhibitors depend on the carboxylic acid functionality to compete with the SF3B1 conformational change required for intron engagement, while inactive competitors are essentially ejected from SF3B upon closing. We aim to test this model in the future with inactive herboxidiene analogs that are engineered with a cross-linking group to stabilize their occupancy in the channel.

**Figure 6.**
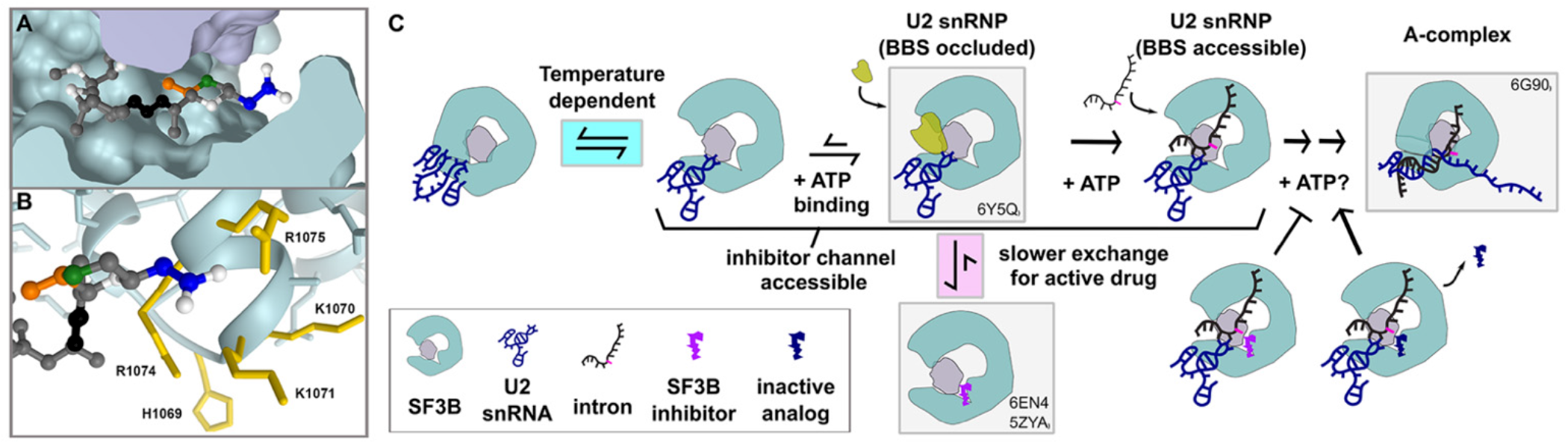
Model of early spliceosome assembly and SF3B inhibition. (a) Model of herboxidiene bound to SF3B inhibitor channel between SF3B (teal) and PHF5A (purple). Positions are colored as follows: C1 blue, C5 green, C6 orange, diene black. (b) Sidechains located near the modeled position of herboxidiene C1. Mutations of labeled residues in yellow confer resistance to Pladienolide B or herboxidiene. C. Model of early spliceosome assembly steps and SF3B inhibitor action. Complexes boxed in gray are supported by indicated structure models ^1,18,25,26^.

Regardless of mechanism, the ability of iHB (**14**) to compete with inhibitors provides unique way to study the parameters that affect their activity. Order of addition experiments revealed the inhibitors behave as if they have a very slow off-rate, because once SF3B is exposed to the compounds, excess competitor loses it advantage relative to when the inhibitor and competitor are added in reverse order. We also see that temperature affects SF3B inhibitor interactions, which we hypothesize relates to the transition between closed and open SF3B1 conformations (Figure 6c). Because colder temperature results in less inhibitor interaction, we predict the closed conformation is stabilized at 4°C relative to 30°C, the temperature at which extracts are most active for splicing. We showed that the effect of temperature is not due to an ATP hydrolysis event, although there is a potential ATP-binding event that further stabilize SF3B1 in the open conformation at any temperature. We also mapped an ATP-dependent rearrangement in U2 snRNP that allows access of the branch binding sequence to base pairing with a complimentary oligo, although this change was independent of SF3B inhibition.

Previous observations show that in the presence of SF3B inhibitors U2 snRNP is able to join an intron in an A-like splicing complex ^21–23^. However, that complex is less stable relative to the real A-complex, and likely has not yet formed the branch helix ^22,29^. Notably even in the presence of inhibitor, ATP-hydrolysis is required for U2 snRNP’s interaction with an intron. We hypothesize that this event may be related to the rearrangement that leads to increased accessibility of the U2 snRNA branch binding sequence. Notably, a recent cryo-EM structure revealed a population of U2 snRNPs in which the protein HTATSF1 is situated between the open “jaws” of SF3B1, while the branch binding sequence of U2 snRNA is folded in the BSL and surrounded by both HTATSF and DDX46 ^18^. It may be that ATP-binding by DDX46 promotes these interactions to help stabilize the SF3B1 open conformation necessary for inhibitor occupancy of the binding channel (Figure 6C). We speculate that an ATP-dependent restructuring of U2 snRNP, potentially involving the removal of HTATSF1 and/or release of DDX46 would allow an intron initial access to the BSL to form the A-like complex stalled by SF3B inhibitors. We also suspect additional ATP-dependent rearrangements in U2 snRNP are required for branch helix formation given the extensive rearrangements of RNA/RNA and RNA protein interactions that must take place for U2 snRNP to take on the structure observed in the cryo-EM model of A-complex^1^. Nearly irreversible inhibitor interaction with SF3B prevents the transition to a closed SF3B1 conformation that is required to form or stabilize the branch helix. Nailing down the players and order of these interactions is the next challenge for splicing researchers, and SF3B inhibitors will likely continue to be important tools for helping to probe the process.

## MATERIALS AND METHODS

### Synthesis of SF3B1 inhibitors and analogs

Structures of SSA, PB, herboxidiene, iSSA, iPB and iHB are shown in Supplemental Figure 4. Synthesis of SF3B1 inhibitors and herboxidiene analogs (**2**), (**3**), (**5**) and (**14**), are published^41,43–46^. Herboxidiene analogs (**4**), (**6-3**) and (**15-20**) were synthesized and purities were assessed by HPLC (>90% pure). The details of synthesis will be published elsewhere (Ghosh et al, *in preparation*).

### *In vitro* splicing analysis

Pre-mRNA substrate was derived from the adenovirus major late (AdMl) transcript (sequence shown in Supplemental Figure 5). ^32^P-UTP body-labeled G(5’)ppp(5’)G-capped substrate was generated by T7 run-off transcription followed by gel purification. Nuclear extract was prepared as previously described ^47^ from Hela cells grown in DMEM/F-12 1:1 and 5% (v/v) newborn calf serum. Order of addition experiments started with 60 mM potassium glutamate, 2 mM magnesium acetate, 2 mM ATP, 5 mM creatine phosphate, 0.05 mg/ml tRNA, 50% (v/v) Hela nuclear extract and first compound and incubated at 4°C or 30 °C. Next the second compound was added, and the mixture was incubated for 10, 30 or 90 minutes at 4°C or 30°C. Then, radiolabeled pre-mRNA was added to a final concentration of 10 nM and the mixture incubated at 30°C for 30 or 60 minutes. The RNA was isolated from the reactions and separated on a 15% (v/v) denaturing polyacrylamide gel. 32P-labeled RNA species were visualized by phosphorimaging and quantified with ImageQuant software (Molecular Dynamics). Splicing efficiency was quantified as the amount of mRNA relative to total RNA and normalized to a dimethyl sulfoxide (DMSO) control reaction.

For experiments testing ATP-dependency, Hela nuclear extract was first depleted of ATP by incubation at 30°C for 15 minutes. Following inhibitor competitions with and without added ATP (2 mM), additional ATP (2 mM) was included with pre-mRNA substrate to allow for splicing.

### RNase H cleavage analysis

A mixture of 40% Hela nuclear extract (depleted of ATP by incubating for 10 minutes at 30°C) 60 mM potassium glutamate, 2 mM magnesium acetate, 0.1 mg /ml tRNA with and without 2 mM ATP and 1μM SSA or DMSO was incubated for 10 minutes at 30°C. DNA oligos complementary to U2 snRNA 24-38nt or 32-46nt were added at 5 μM and incubated for another 15 minutes at 30°C to allow for cleavage by endogenous RNase H. The oligos were degraded with addition of 1 μl RQ1 DNase for 10 minutes at 30°C. RNA was isolated and used as template for reverse-transcription primer extension reactions by first annealing with a 32P-radiolabeled primer complementary to U2 snRNA nt 97-117. Reverse transcription reactions contained 50 mM Tris pH 7.9, 75 mM potassium chloride, 7 mM DTT, 3 mM magnesium chloride, 1 mM dNTPs, and 0.5 μg reverse transcriptase (MMlV variant). After a 30 minute incubation at 53°C, DNA was isolated and separated on a 10% (v/v) denaturing polyacrylamide gel that was dried and visualized as described previously.

## ACKNOWLEDGEMENTS

This work was funded by National Institute of General Medical Sciences (NIGMS) of the National Institutes of Health (NIH) under award R01GM72649 to M.S.J and A.K.G. A.G.l. was supported by NIGMS award R25GM058903 and by the National Science Foundation Graduate Research Fellowship under grant DGE-1842400

