## Supplemental Figures for "Identification of herboxidiene features that mediate conformation-dependent SF3B1 interactions to inhibit splicing"

**SUPPLEMENTAL DATA**

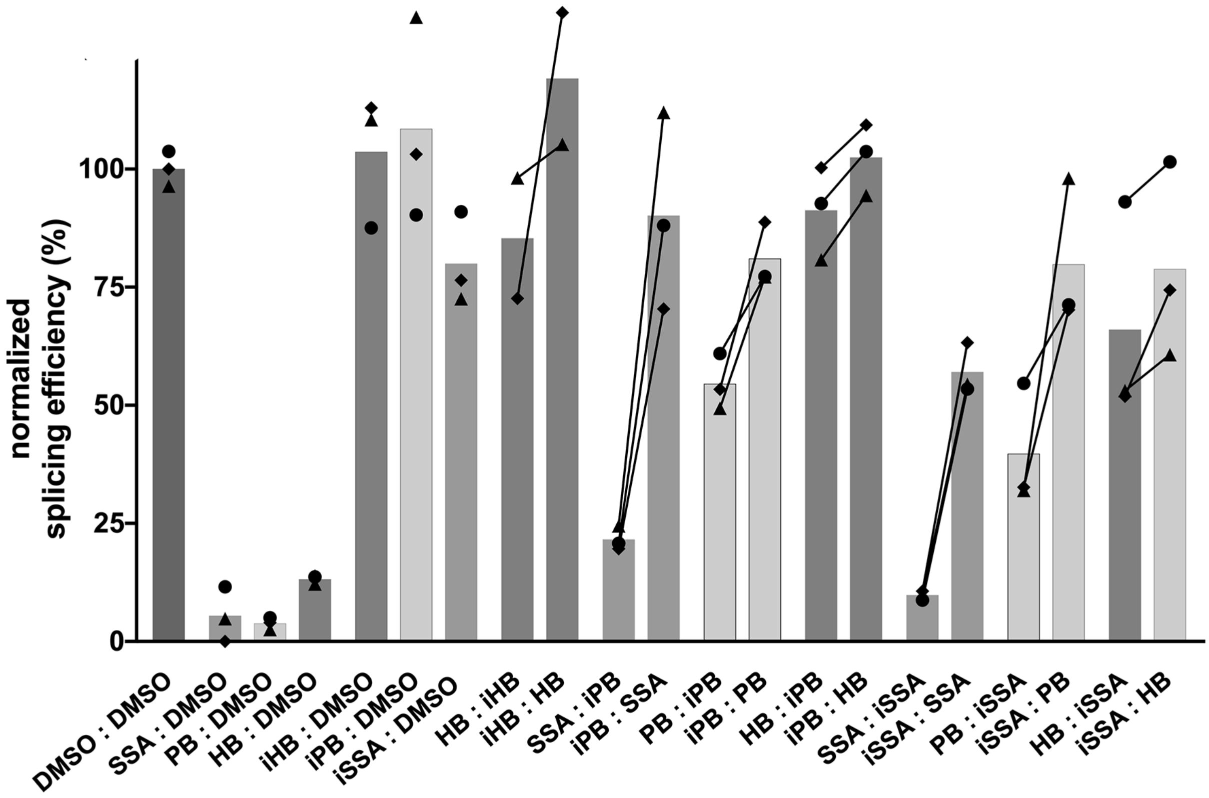


**Supplemental Figure 1: Order of addition affects competition between SF3B inhibitors and inactive PB and SSA analogs**

Quantification of in vitro splicing assays. Prior to addition of splicing substrate, nuclear extract was incubated at 4°C for 10 minutes with one of the compounds DMSO, herboxidiene (HB, 1µM), SSA (1µM) PB (1µM) iHB (100µM), iSSA (100 µM) or iPB (100 µM) followed by the indicated compound at same concentrations for another 10 minutes. Mean normalized splicing efficiency is displayed as bars, with experimental replicates values represented by different shape and linked to illustrate the difference in splicing rescue.

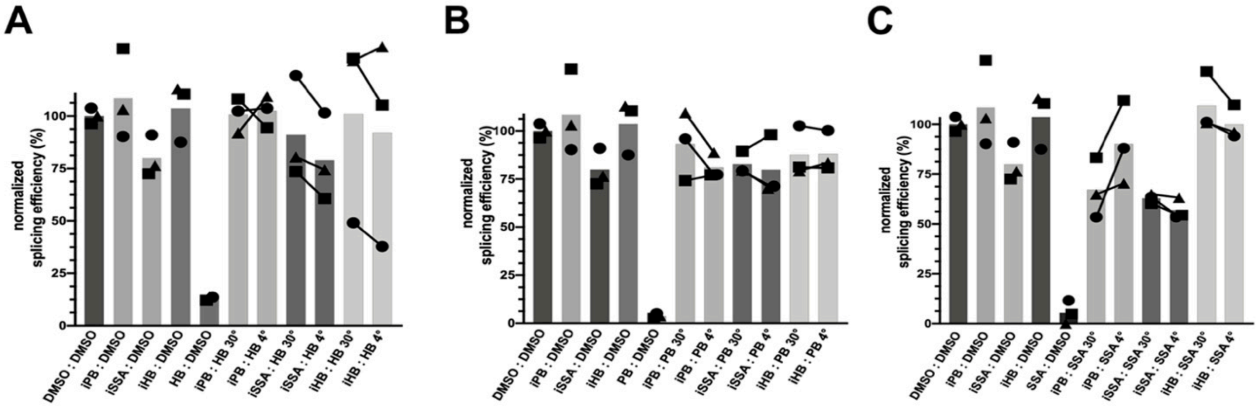


**Supplemental Figure 3: Temperature dependent inhibition of splicing depends on order of addition of excess inactive competitor**

Quantification of *in vitro* splicing assays. Prior to addition of splicing substrate, nuclear extract was incubated at 4°C or 30°C for 10 minutes with one of the compounds DMSO, herboxidiene (HB, 1µM), SSA (1µM) PB (1µM) iHB (100µM), iSSA (100 µM) or iPB (100 µM) followed by addition of DMSO, herboxidiene (HB, 1 µM; panel A), SSA(1 µM; panel B) or iPB (100 µM; panel C) at the same concentration and temperature for another 10 minutes. Mean normalized splicing efficiency is displayed as bars, with experimental replicates values represented by different shape and linked to illustrate the difference in splicing rescue.

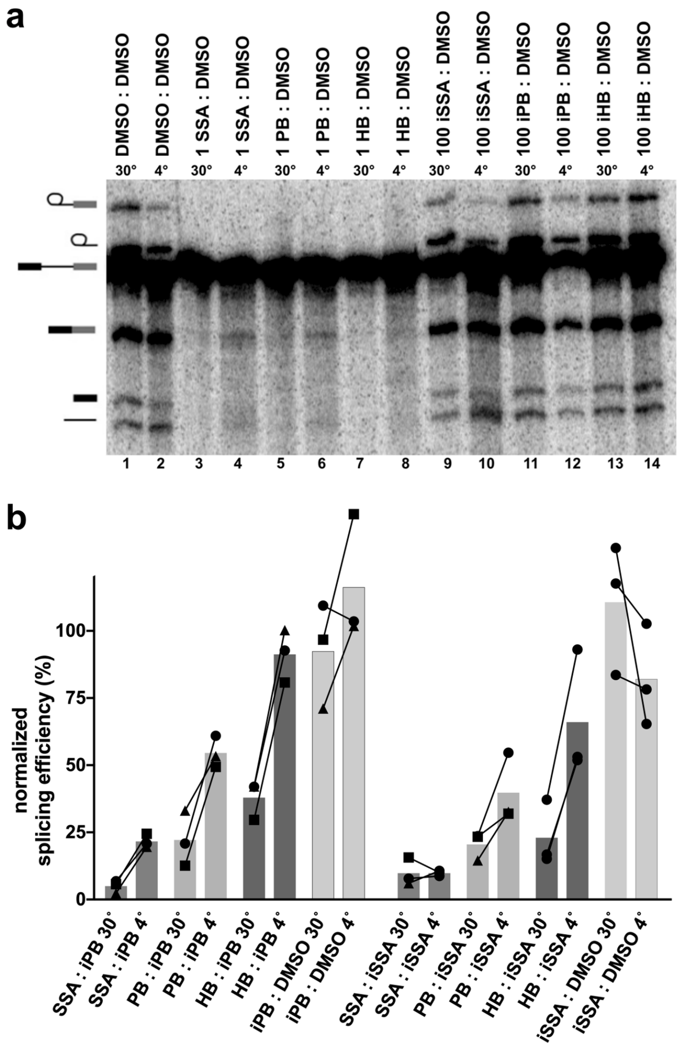


**Supplemental Figure 2: Temperature affects competition with iSSA and iPB but not inactive and active compound controls**

(a) Representative denaturing gel analysis of *in vitro* splicing using nuclear extracts incubated at 4˚C or 30˚C for 10 minutes with DMSO, 1 µM (SSA, PB or HB) or 100 µM (iSSA, iPB,iHB), followed by DMSO for 10 minutes. Band identities are illustrated on the left as (from top to bottom) lariat intermediate, lariat intron product, pre-mRNA substrate, mRNA product, 5' exon intermediate and linear intron product. (b) Quantification of *in vitro* splicing assays. Prior to addition of splicing substrate, nuclear extract was incubated at 4°C or 30°C for 10 minutes with one of the compounds DMSO, herboxidiene (HB, 1µM), SSA (1µM) PB (1µM), iSSA (100 µM) or iPB (100 µM) followed by addition of DMSO, iPB (100 µM) or iSSA (100 µM B) at the same concentration and temperature for another 10 minutes. Mean normalized splicing efficiency is displayed as bars, with experimental replicates values represented by different shape and linked to illustrate the difference in splicing rescue.

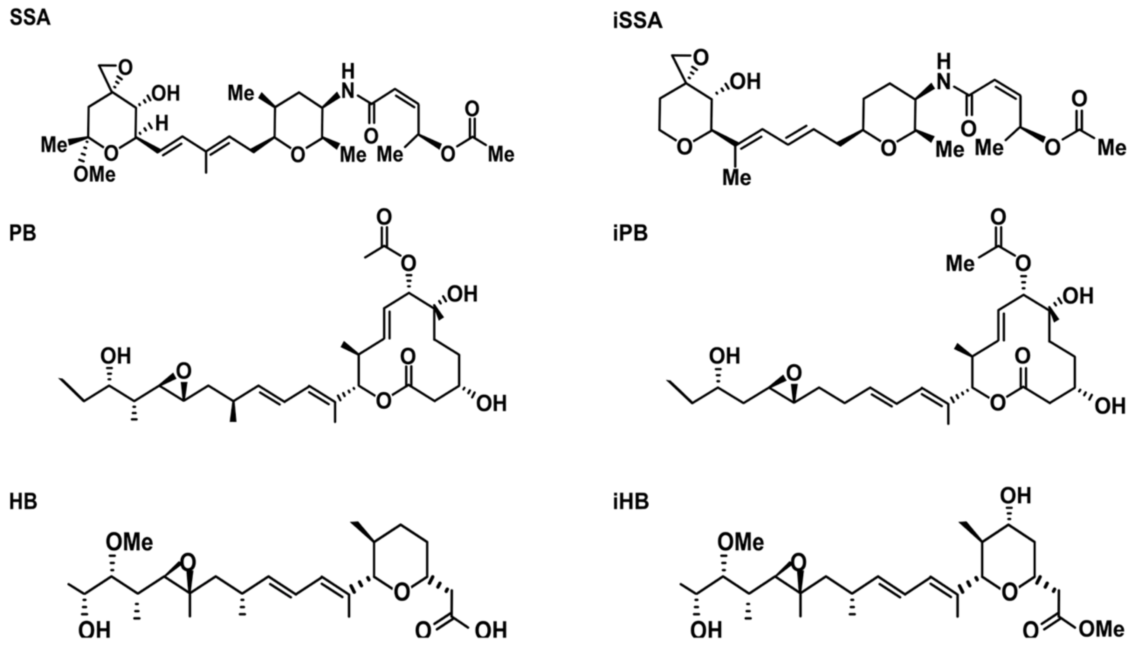


**Supplemental Figure 4: Chemical structures of SF3B inhibitor analogs used in this study**

GAGACCGGCAGATCAGCTTGGCCGCGTCCATCTGGTCATCTAGGATCTGATATCATCGATGA
ATTCGAGCTCGGTACCCCGTTCGTCCTCACTCTCTTCCGCATCGCTGTCTGCGAGGGCCAGCG
TAAAAGGTGAGTACTCCCTCTCAAAAGCGGGCATGACTTCTGCCCTCGAGTTATTAACCCTC
ACTAAAGGCAGTAGTCAAGGGTTTCCTTGAAGCTTTCGTGCTGACCCTGTCCCTTTTTTTTCC
ACAGCTGCAGGTCGACGTTGAGGACAAACTCTTCGCGGTCTTTCCAGTACTCTTG

**Supplemental Figure 5: Pre-mRNA sequence of substrate used in splicing for this study**
